# Hyperglycemia and Cancer. Human lung carcinoma by means of Raman spectroscopy and imaging

**DOI:** 10.1101/2022.04.05.487128

**Authors:** H. Abramczyk, M. Kopeć, K. Beton

## Abstract

Raman spectroscopy and Raman imaging allow to identify the biochemical and structural features of human cancer lung cell line (CCL-185) and the cell line supplemented with glucose and deuterated glucose in normal and hyperglycemia conditions. We found that isotope substitution of glucose by deuterated glucose allows to separate *de novo* lipid synthesis from exogenous uptake of lipids obtained from the diet. We demonstrated that glucose is largely utilized for *de novo* lipid synthesis. Our results provide a direct evidence that high level of glucose decreases the metabolism via oxidative phosporylation in mitochondria in cancer cells and shifts the metabolism to glycolysis via Wartburg effect.

**Graphical Abstract:** 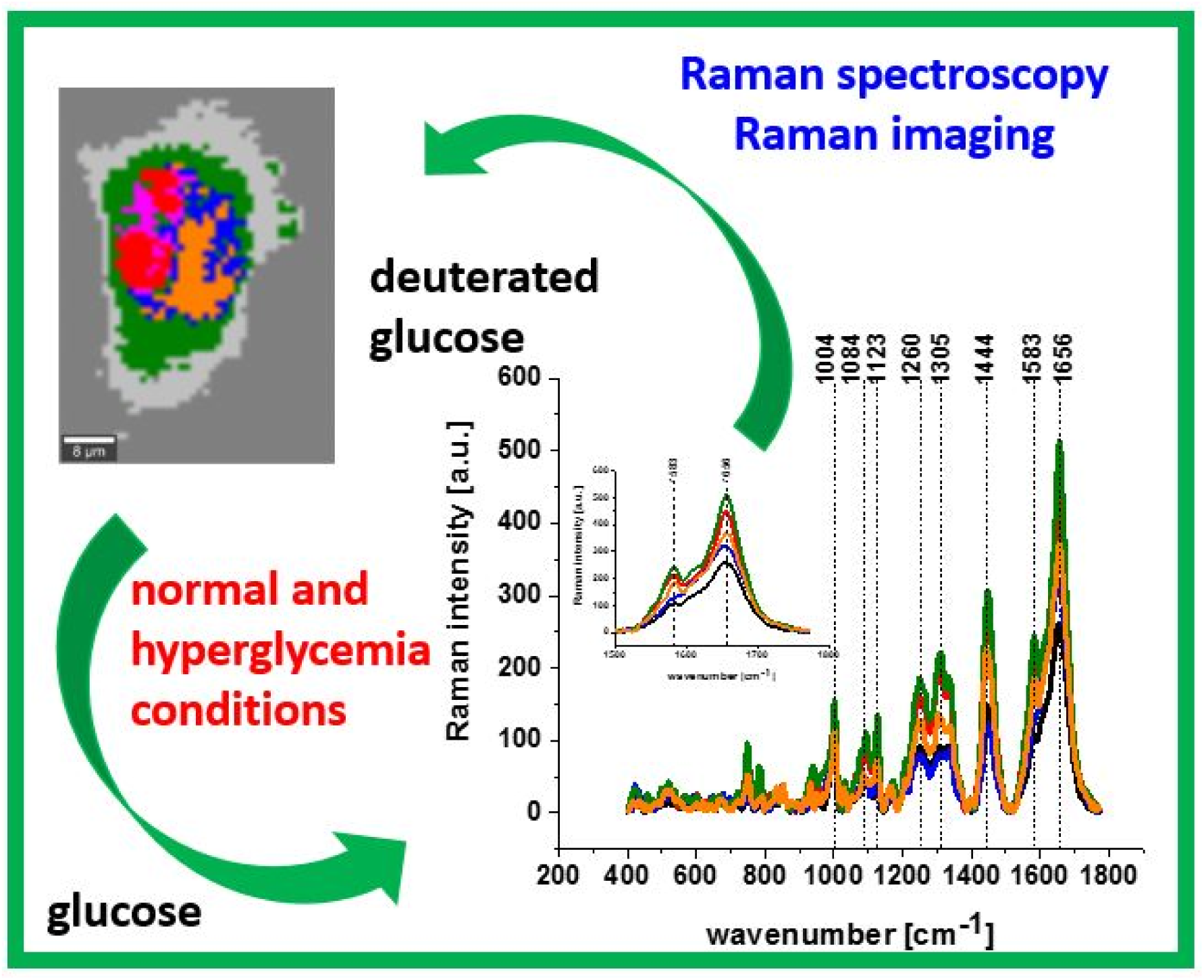

## Introduction

Sugars are molecules contain carbon, hydrogen and oxygen atoms. Sugars play a lot of important role in the human body. First of all sugars are source of energy for human body. Another important function of sugars is fueling the body. [1] Moreover sugars are one of the most important regulators of physiological processes in body such as growth, stress responses.

Diabetes is defined as a metabolic disease characterized by elevated levels of blood sugar, which leads to damage to many of the body’s systems. According to World Health Organization diabetes is called 21^st^ century epidemic. [2] It is estimated that the number of obese will be increase more rapidly in developing regions of the world. [3]

The values for normal fasting blood glucose concentration are between 3.9-5.6 mmol/L. The expected values for normal fasting blood glucose concentration are between 70 mg/dL (3.9 mmol/L) and 100 mg/dL (5.6 mmol/L). When fasting blood glucose is between 100 to 125 mg/dL (5.6 to 6.9 mmol/L) changes in lifestyle and monitoring glycemia are recommended.

Isotope substitution has been used to study metabolic processes and their molecular mechanisms of individual living cells. [4] [5] [6] [7] [8] [9] [10] [11] [12] [13] [14] [15] [16] [17] [18] [19] [20]

Various existing techniques of molecular biology methods such as enzyme-linked immunosorbent assays (ELISA), Western blot, high performance liquid chromatography (HPLC), spectrophotometry and flow cytometry can be used to estimate concentration of metabolites. Isotope substitution method has been extensively used in mass spectrometry (MS), nuclear magnetic resonance spectroscopy and Raman spectroscopy to identify and quantify metabolites with high sensitivity. [7] [4] [5] [8] [9–13] [14] [15] However, the implementation of these techniques is limited due to their destructive nature (MS) or inability to detect the full extent of metabolite localization inside and outside specific organelles.

Therefore, these methods provide only bulk analyses of cell populations and cannot monitor the biochemical heterogeneity within specific organelles of a single cell. Moreover, none of the methods used to control metabolites concentration can provide direct evidence about their role in apoptosis and oxidative phosphorylation, because they are not able to monitor the amount of concentration in specific organelles such as mitochondria, cytoplasm, or extracellular matrix. Therefore, existing analytical technologies cannot detect the full extent of specific metabolites or their biomarkers localization inside and outside specific organelles. In Raman imaging we do not need to disrupt cells to release the cellular structures to learn about their biochemical composition. [21] [22] [23] [24] [25] [26] [27] Recently isotopic substitution has been used in linear and nonlinear Raman imaging. [28] [29] [30]

Given that hyperglycemia is the most important biological feature of diabetes and cancer needs lots of glucose, it may suggest that hyperglycemia plays an important role during cancer progression in cancer. [27] Increasing evidence suggests a close association between diabetes and cancer; however, the links between the two diseases remain unclear. [31]

In this study, we applied Raman imaging to study metabolism of human cancer lung cell line (CCL-185) and the cell line supplemented with glucose and deuterated glucose in normal and hyperglycemia conditions. We will concentrate on *de novo* lipogenesis, which is being increasingly recognized as a hallmark of cancer. [32] [33] [34]

The aim of this paper is to demonstrate the possibility of monitoring glucose changes occurring in cancer cells by Raman spectroscopy and Raman imaging. The aim of this work is to understand the role of sugars in metabolism of human lung cancer cell lines CCL-185. In this paper we have analyzed the biochemical composition of lung cancer cell lines CCL-185 and cell lines supplemented with glucose and deuterated glucose. The experiments were carried out using Raman spectroscopy and Raman imaging.

Currently used therapies don’t perfectly address many type of diabetes. [35] It is very justified to develop new methods to complete this gap. Raman methods give fast, objective information about biochemical composition in specific organelles of single cells. Understanding sugar’s role in lung cancer with using Raman spectroscopy will help establish a modern clinical diagnosis tools for diabetes monitoring.

## Materials and methods

### Chemicals

D-(+)-glucose (product number G7021) and D-glucose-1,2,3,4,5,6,6-d_7_ (product number 552003) were purchased from MERCK.

### Raman spectroscopy

All Raman spectra and Raman images presented in this manuscript were recorded using WITec alpha 300 RSA + combined with confocal microscope coupled via the fibre of a 50 μm core diameter with an UHTS (Ultra High Throughput Spectrometer) spectrometer and a CCD Camera (Andor Newton DU970N-UVB-353) operating in standard mode with 1600×200 pixels at −60 °C with full vertical binning. All experiments were performed using laser diode (SHG of the Nd:YAG laser (532 nm)) and 40x Nikon water dipping objective (NA = 1.0). All cells were measured using laser with a power 10 mW at the sample position. The Raman images for all cells were recorded with integration time 0.3 sec in the high frequency region and with 0.5 sec in the fingerprint region. To acquisition and preprocess of the data (cosmic rays removing, smoothing and removing background) we used WITec Project Plus. All Raman maps were analyzed by Cluster Analysis method. [36] [37]

Spectroscopic data were analyzed using Cluster Analysis method. Briefly, Cluster Analysis is a form of exploratory data analysis in which observations are divided into different groups that have some common characteristics – vibrational features in our case. Cluster Analysis constructs groups (or classes or clusters) based on the principle that within a group of observations (the Raman spectra) must be as similar as possible, while the observations (the Raman spectra) belonging to different groups must be different. The partition of n observations (x) into k (k≤n) clusters S should be done to minimize the variance (Var) according to the formula:

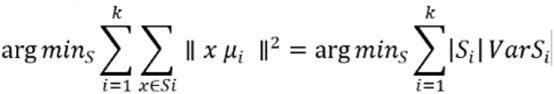

Where *μi* is the mean of points The Raman maps presented in the paper were constructed based on the principles of Cluster Analysis described above. The number of clusters was 7. Each cluster is characterized by specific average Raman spectra, which reflects the inhomogeneous distribution of chemical components within the organelles of single cells.

### Cell lines and Cell Culture

A549 cell line (ATCC® CCL-185™) was purchased from ATCC: The Global Bioresource Center. CCL-185 cell line was cultured using ATCC-formulated F-12K Medium (Kaighn’s Modification of Ham’s F-12 Medium), Catalog No. 30-2004, contains 2 mM L-glutamine and 1500 mg/L sodium bicarbonate. To make the complete growth medium, Fetal Bovine Serum (FBS) was added to a final concentration of 10%. The culture medium was renewed from 2 to 3 times a week. The CCL-185 cells obtained from the patient are isolated from the lung tissue of a 58-year-old Caucasian male with lung cancer. The biological safety of the CCL-185 cell line has been classified by the American Biosafety Association (ABSA) as level 1 (BSL-1).

### Cultivation Conditions

The cell line (CCL-185) used in the experiments in this study were grown in flat-bottom culture flasks made of polystyrene with a cell growth surface of 75 cm^2^. Flasks containing cells were stored in an incubator providing environmental conditions at 37 °C, 5% CO2, 95% air.

### Cell Treatment with Glucose and Deuterated glucose

Cells used for research were seeded onto CaF_2_ windows (25 × 1 mm) at a low density of 104 cells/cm^2^. After 24 h incubation on the CaF2, standard growth medium was removed, and glucose or deuterated glucose in concentrations simulating normal (5 mM) and hyperglycemic (100 mM) conditions solution was added for 24 h. After 24h, the cells were rinsed with Phosphate Buffered Saline (PBS, Gibco, 10010023, pH 7.4 at 25°C, 0.01 M) to remove any residual medium and an excess of additives that did not penetrate inside the cells. Furthermore, PBS was removed, and cells were fixed in Formaldehyde (4% buffered formalin) for 10 min and washed once more with PBS. The Raman confocal measurements were made immediately after the preparation of the samples. All the glucose solutions used for supplementation procedure in investigation were prepared by diluting in the pure culture medium dedicated to cell lines used in investigation as a solvent. For the preparation of glucose and deuterated glucose solutions, powdered reagents were used, from which samples were prepared to finally obtain a solution with concentrations of 5 and 100 mM.

## Results and discussion

First, we focused on the Raman spectroscopy and Raman imaging analysis for lung cancer cells CCL-185 without any glucose supplementation. Figure 1 presents video image, Raman images and Raman spectra for high frequency region and for low frequency region for CCL-185 human lung cancer cell.

**Fig. 1.**
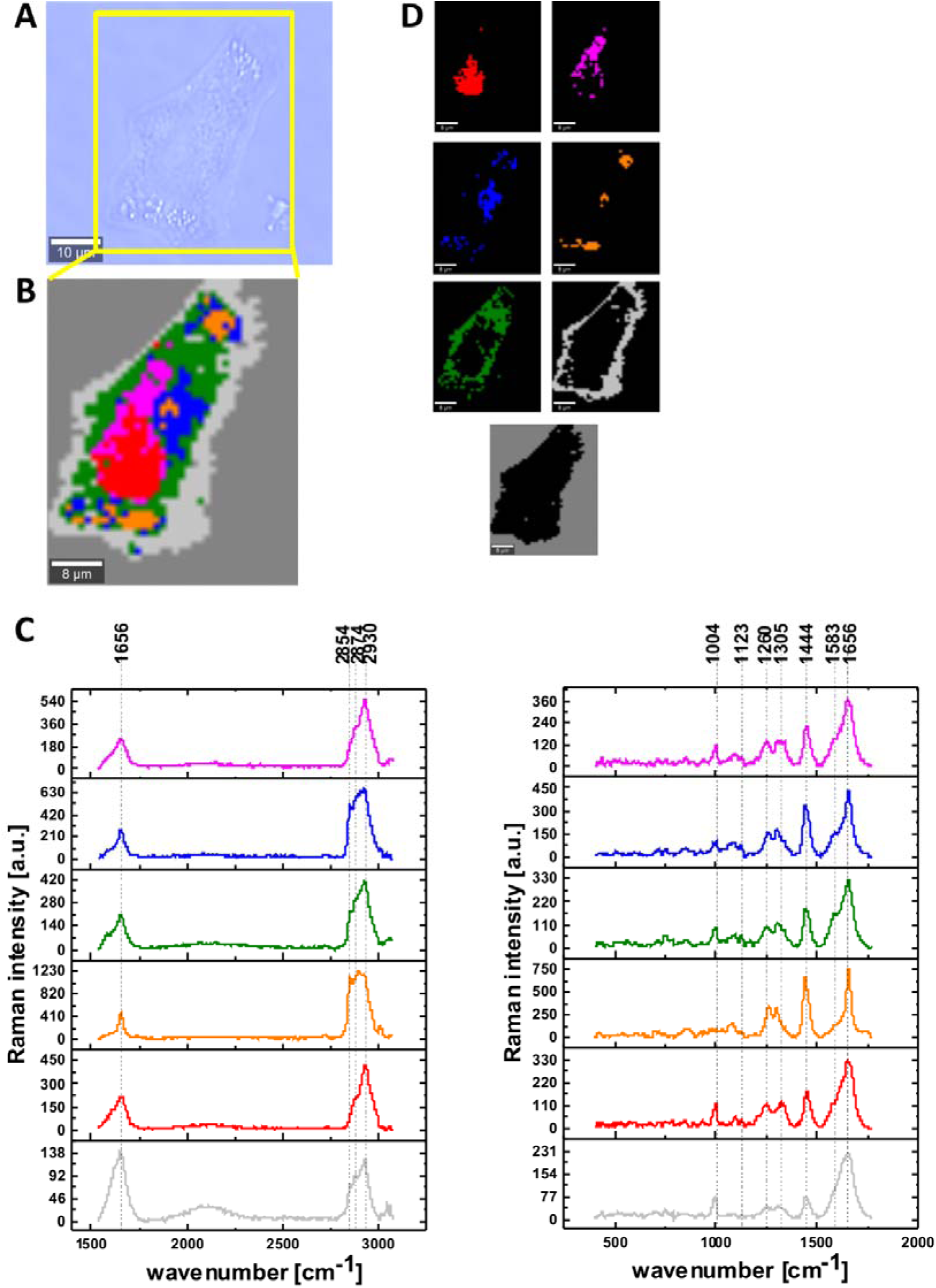
The microscopy image (A), Raman image for area marked by yellow frame in the panel A, the size of Raman image (45 μm × 42 μm), resolution 1 μm of human lung single cell CCL-185 constructed based on Cluster Analysis method (B), the average Raman spectra for all clusters for high and for low frequency region (C), Raman images of separate clusters identified by Cluster Analysis method assigned to: nucleus (red), mitochondria (magenta), lipid regions (blue and orange), cytoplasm (green), cell membrane (light grey) and cell environment (dark grey) (D), colors of the spectra correspond to the colors of clusters; integration time 0.3 sec in the high frequency region and 0.5 sec in the fingerprint region, laser power 10mW.

To define the main biochemical components in single cells we used Cluster Analysis method. In Fig. 1 panel D we present clusters identified by Cluster Analysis. The red area shows the locations of nucleus, magenta areas shows the locations of mitochondria, blue and orange areas reflect the locations of lipids in rough lipid droplets and endoplasmic reticulum respectively, green areas reflect the locations of cytoplasm, light gray areas reflect the locations of cell membrane and dark gray areas reflect the locations of cell environment. One can see from Fig.1 (panels A, B, D) an ideal match between Raman maps and microscopy image.

The results presented in Fig. 1C confirm that Raman imaging can characterize the biochemical composition of organelles in human lung cells. One can see that the single cell in fingerprint region is dominated by peaks at: 1004, 1123, 1260, 1305, 1444, 1583, 1656 cm^-1^ and by peaks at: 2854, 2874, 2930 cm^-1^ in the high frequency region. Table 1 presents the main chemical components which can be identified based on their vibrational features.

**Table 1.**
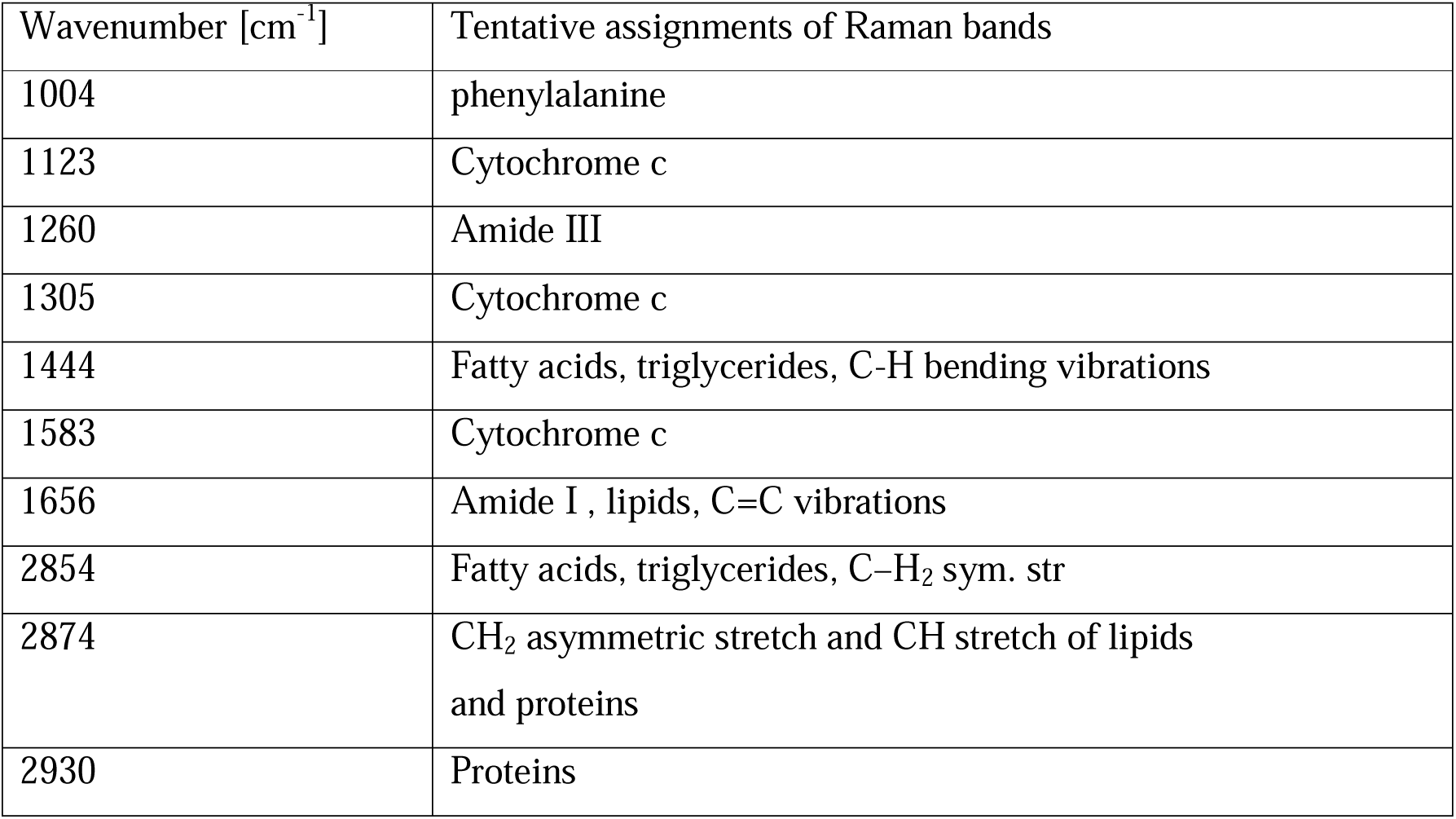
Band positions for human lung cells [38] [39] [32]

| Wavenumber [$\text{cm}^{-1}$] | Tentative assignments of Raman bands |
| --- | --- |
| 1004 | phenylalanine |
| 1123 | Cytochrome c |
| 1260 | Amide III |
| 1305 | Cytochrome c |
| 1444 | Fatty acids, triglycerides, C-H bending vibrations |
| 1583 | Cytochrome c |
| 1656 | Amide I, lipids, C=C vibrations |
| 2854 | Fatty acids, triglycerides, C-H<sub>2</sub> sym. str |
| 2874 | CH<sub>2</sub> asymmetric stretch and CH stretch of lipids and proteins |
| 2930 | Proteins |

As we mentioned above the main aim of our paper was to investigate the changes in cell metabolism after supplementation with glucose at normal and hyperglycemia conditions. Fig. 2 presents Raman spectra for glucose and deuterated glucose in PBS.

**Fig. 2.**
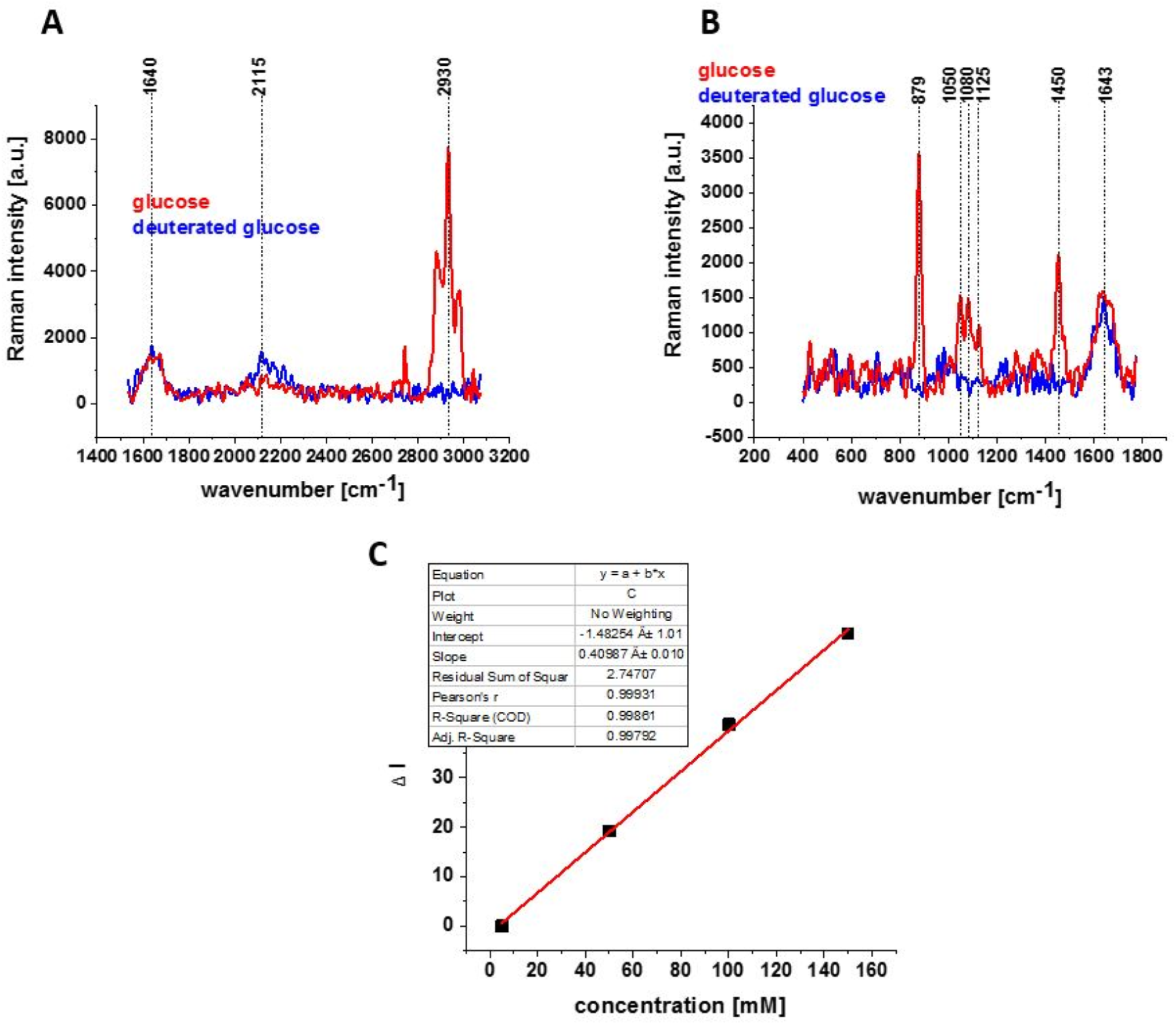
Raman spectra for glucose and deuterated glucose d-7 (150 mM) in PBS in high frequency region (A) and in low frequency region (B); integration time 20 s, 1 accumulation, laser power 10 mW; Linear dependence (of Raman intensity of the band at 2120 cm^-1^-background) on glucose d-7 concentration; integration time 1 s, 10 accumulation, laser power 10 mW (C)

Comparison of the Raman spectrum for deuterated and normal glucose shows several important difference, which will used in our studies. First, peaks at around 2930 cm^-1^ are very intense in Raman spectrum for glucose (red line). They correspond to C-H stretching vibrations of the CH_2_ or CH_3_ groups. Each D substitution in CH_2_ or CH_3_ groups of the backbone produces to the band in the C−D stretch region which is shifted to ∼2000−2300 cm^−1^ in the Raman spectrum of deuterated glucose d-7 as predicted by the √2 rule arising from reduced mass effects in the harmonic oscillator model.

One can see that Raman spectrum for deuterated glucose d-7 (blue line) has characteristic wide peak at 2115 cm^-1^. In the fingerprint region one can see strong peaks at 879 cm^-1^ in Raman spectrum for glucose. Fourth, in Raman spectrum for glucose the peak at 1450 cm^-1^ is clearly visible.

The results of Raman spectroscopy and Raman imaging for single cell CCL-185 in hyperglycemia conditions are presented in Fig. 3.

**Fig. 3.**
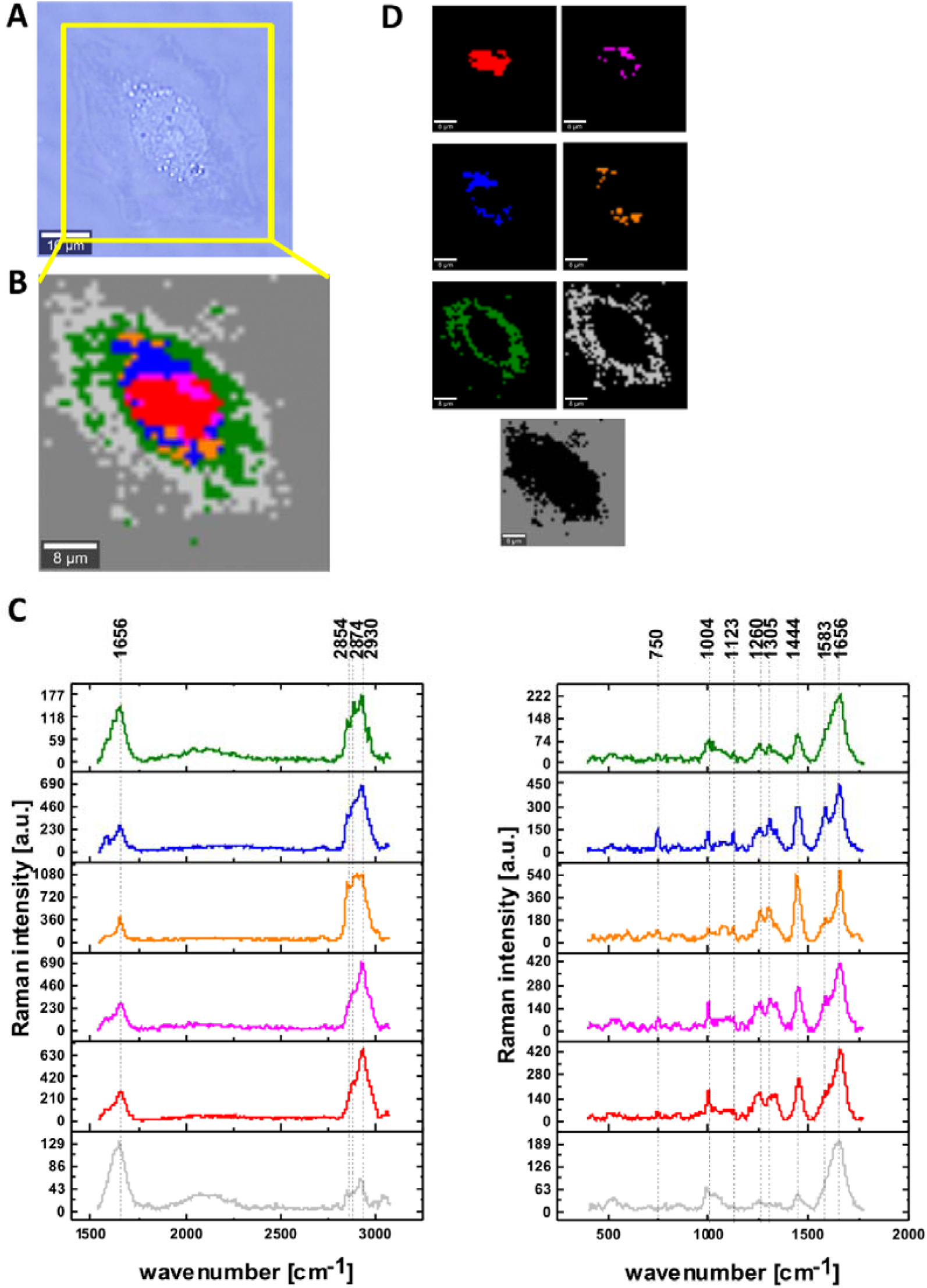
The microscopy image (A), Raman image for area marked by yellow frame in the panel A, the size of Raman image (44 μm × 45 μm), resolution 1 μm of human lung single cell CCL-185 supplemented with glucose 100 mM constructed based on Cluster Analysis method (B), the average Raman spectra for all clusters for high and for low frequency region (C), Raman images of separate clusters identified by Cluster Analysis method assigned to: nucleus (red), mitochondria (magenta), lipid regions (blue and orange), cytoplasm (green), cell membrane (light grey) and cell environment (dark grey) (D), colors of the spectra correspond to the colors of clusters; integration time 0.3 sec in the high frequency region and 0.5 sec in the fingerprint region, laser power 10mW.

In the next step Raman imaging was used to quantify intracellular deuterium content originating from deuterated carbon sources. Deuterium and hydrogen are chemically identical, as deuteration has negligible effect on atom size, molecular shape, equilibrium bond length, or stiffness. [40]

Apart from vibrational effects presented in Fig. 2. We used Raman imaging with isotope labeled glucose at hyperglycemia (100mM) and normal (5mM) conditions to trace the metabolism of glucose with high spatial-temporal resolution.

Figure 4 presents the video image, Raman images and Raman spectra for single human lung cell CCL-185 with 100 mM deuterated glucose at hyperglycemia conditions.

**Fig. 4.**
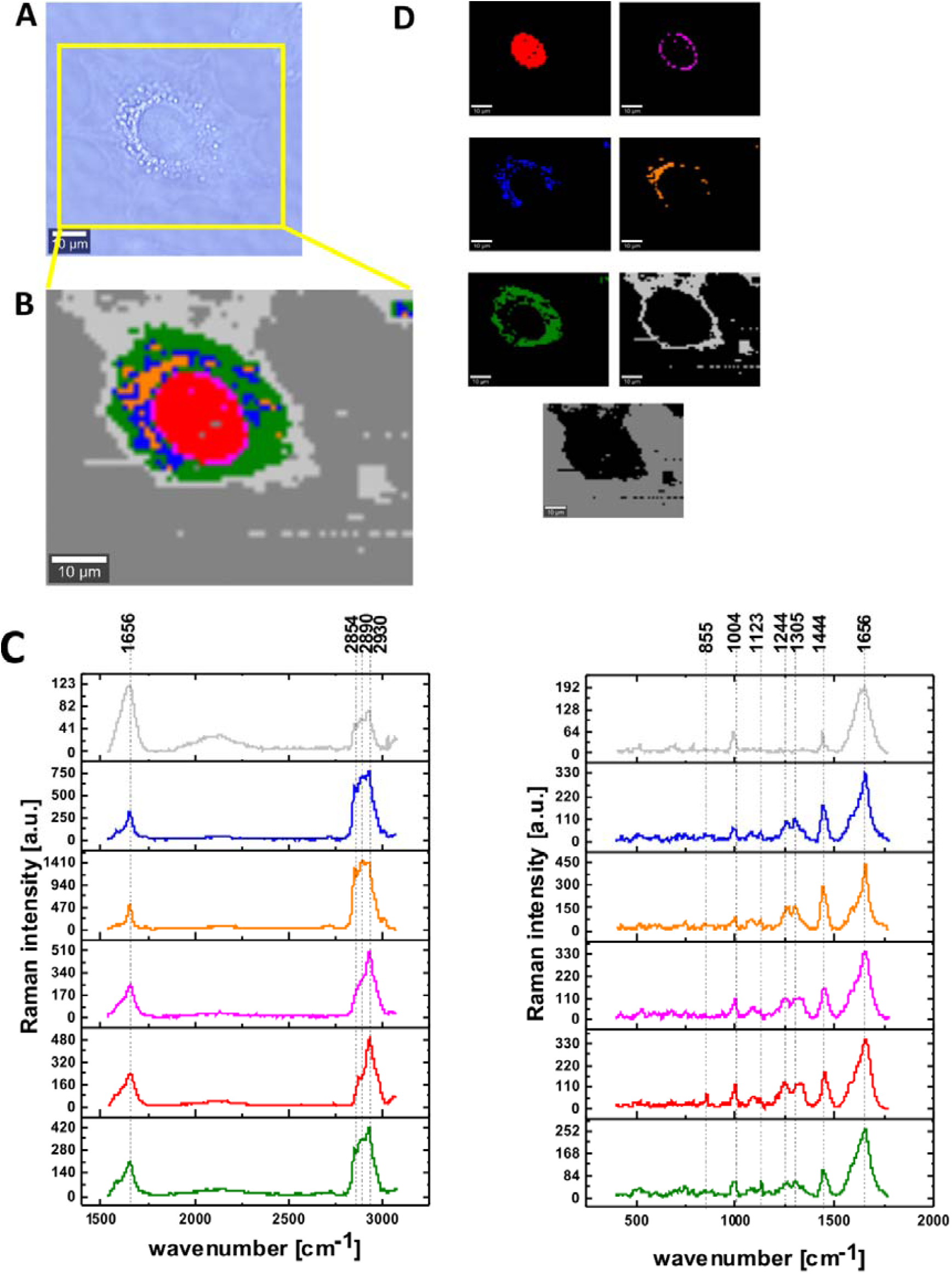
The microscopy image (A), Raman image for area marked by yellow frame in the panel A, the size of Raman image (67 μm × 54 μm), resolution 1 μm of human lung single cell CCL-185 supplemented with deuterated glucose 100 mM constructed based on Cluster Analysis method (B), the average Raman spectra for all clusters for high and for low frequency region (C), Raman images of separate clusters identified by Cluster Analysis method assigned to: nucleus (red), mitochondria (magenta), lipid regions (blue and orange), cytoplasm (green), cell membrane (light grey) and cell environment (dark grey) (D) colors of the spectra correspond to the colors of clusters; integration time 0.3 sec in the high frequency region and 0.5 sec in the fingerprint region, laser power 10mW.

Raman images and Raman spectra for single human lung cell CCL-185 with 5 mM normal and deuterated glucose-d7 at normal conditions are presented in Supplementary Materials (Figures SM1 and SM2).

The main goal of the experiment presented in this study was the biochemical analysis of human lung cell line without supplementation and in normal and hyperglycemia conditions based on the Raman spectra of CCL-185 cell line using Raman spectroscopy and Raman imaging.

To learn more about changes in metabolism we compared the Raman spectra without glucose supplementation, supplemented cells with 5 mM glucose and glucose-d7 (normal conditions) as well as supplemented with 100 mM glucose and glucose-d7 (hyperglycemia conditions). A deuterated carbon atoms in glucose can be used as a universal indicator to probe carbon utilization in glucose metabolism. It is clear from Figures 2-4 that we do not detect glucose or glucose d-7 (lack of the characteristic bands of glucose in the fingerprint region) due to different catabolic pathways degrading glucose. Generally, any possible C−D vibrations of lipids, proteins, nucleic acids, and carbohydrates can give contribution to this spectra region. As lipid biosynthesis contributes roughly 20−35% to the total cellular C−H content, [4] it plays a crucial part in the appearance of the broad C−D band through different lipid catabolic pathways degrading glucose.

In order to better understand the changes caused in metabolism by glucose and deuterated glucose we used the results presented in Figures 1, 3-4 and SM1, SM2 to compare the average Raman spectra for nucleus, mitochondria, cytoplasm, membrane, lipid droplets and endoplasmic reticulum (Fig 5).

**Fig. 5.**
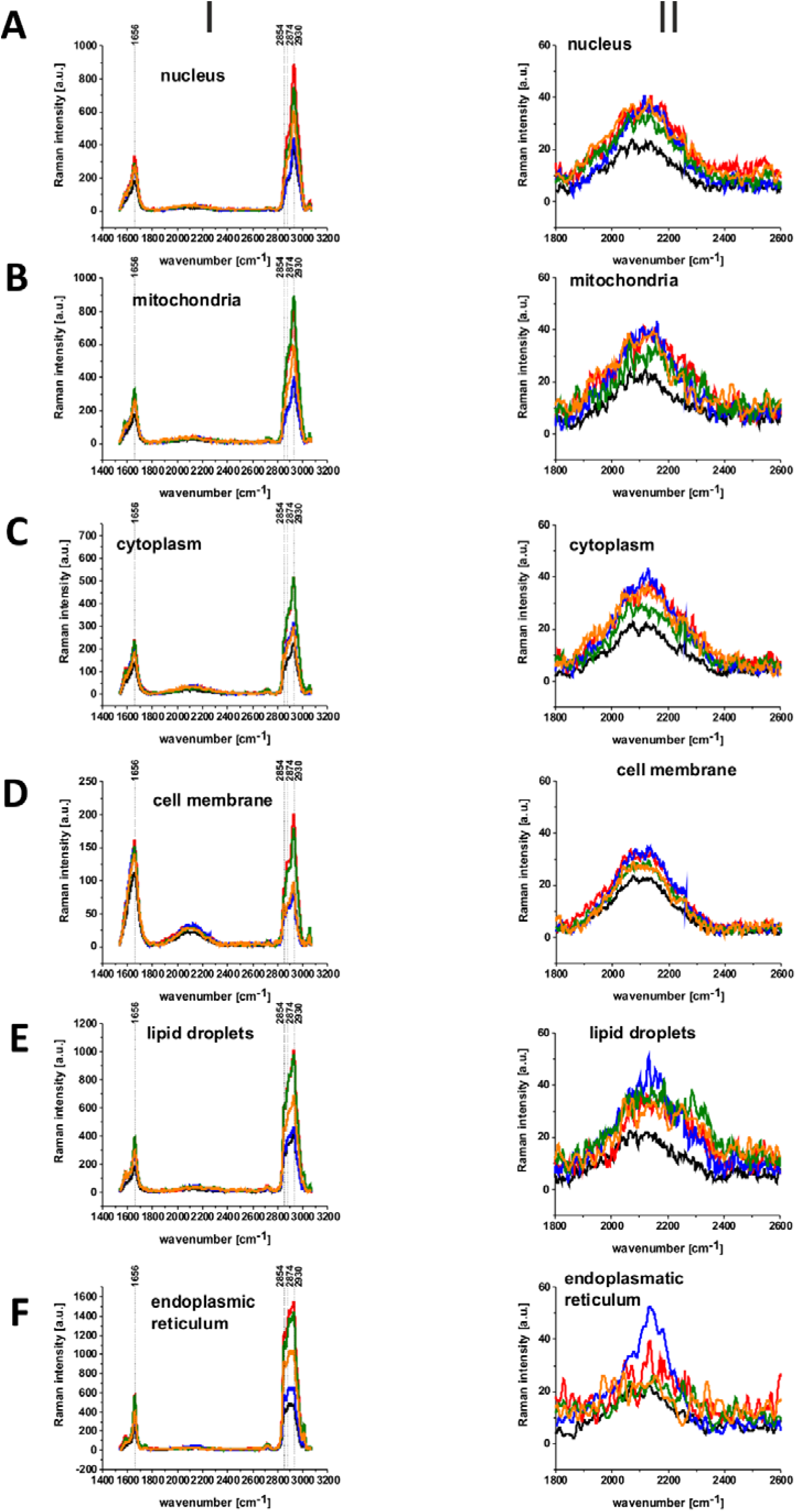
The average Raman spectra of human lung single cell CCL-185 without glucose supplementation (black line), supplemented with normal glucose (green line) and deuterated glucose (red line) at normal conditions (5mM) and supplemented with normal glucose (orange line) and deuterated glucose (blue line) at hyperglycemia conditions (100 mM) for 1400-3200 cm^-1^ spectral region (I) and for 1800-2600 cm^-1^ spectral region (II) obtained from Raman images of separate clusters identified by Cluster Analysis method assigned to: nucleus (A), mitochondria (B), cytoplasm (C), cell membrane (D), lipid droplets (E) and endoplasmic reticulum (F); integration time 0.3 sec in the high frequency region, laser power 10mW.

Here, by coupling Raman imaging with isotope labeled glucose, we were able to trace the metabolism of glucose in single living cells with high spatial-temporal resolution.

First, we compared the Raman signals in the region of C-H stretching vibrations of the CH_2_ or CH_3_ groups of proteins and lipids in the range of 2800-3000 cm^-1^ for supplementation with nondeuterated glucose at normal and hyperglycemia conditions at the same concentration (10%) of lipids from the diet (exogenous uptake in the cellular pool provided by FBS in the growing medium). One can see from Figure 5 that for all organelles the Raman signals corresponding to lipids and proteins are lower at hyperglycemia (100 mM of glucose) than for normal level of glucose (5mM). It indicates that *de novo* lipid synthesis as well as production of proteins decreases at hyperglycemia conditions for cancer lung cells.

It is well known that a common feature of cancer cells is their ability to reprogram their metabolism to sustain the production of ATP and macromolecules needed for cell growth, division and survival [33] [41]

Metabolic pathway in normal cells is presented in the scheme 1.

**Scheme 1.**
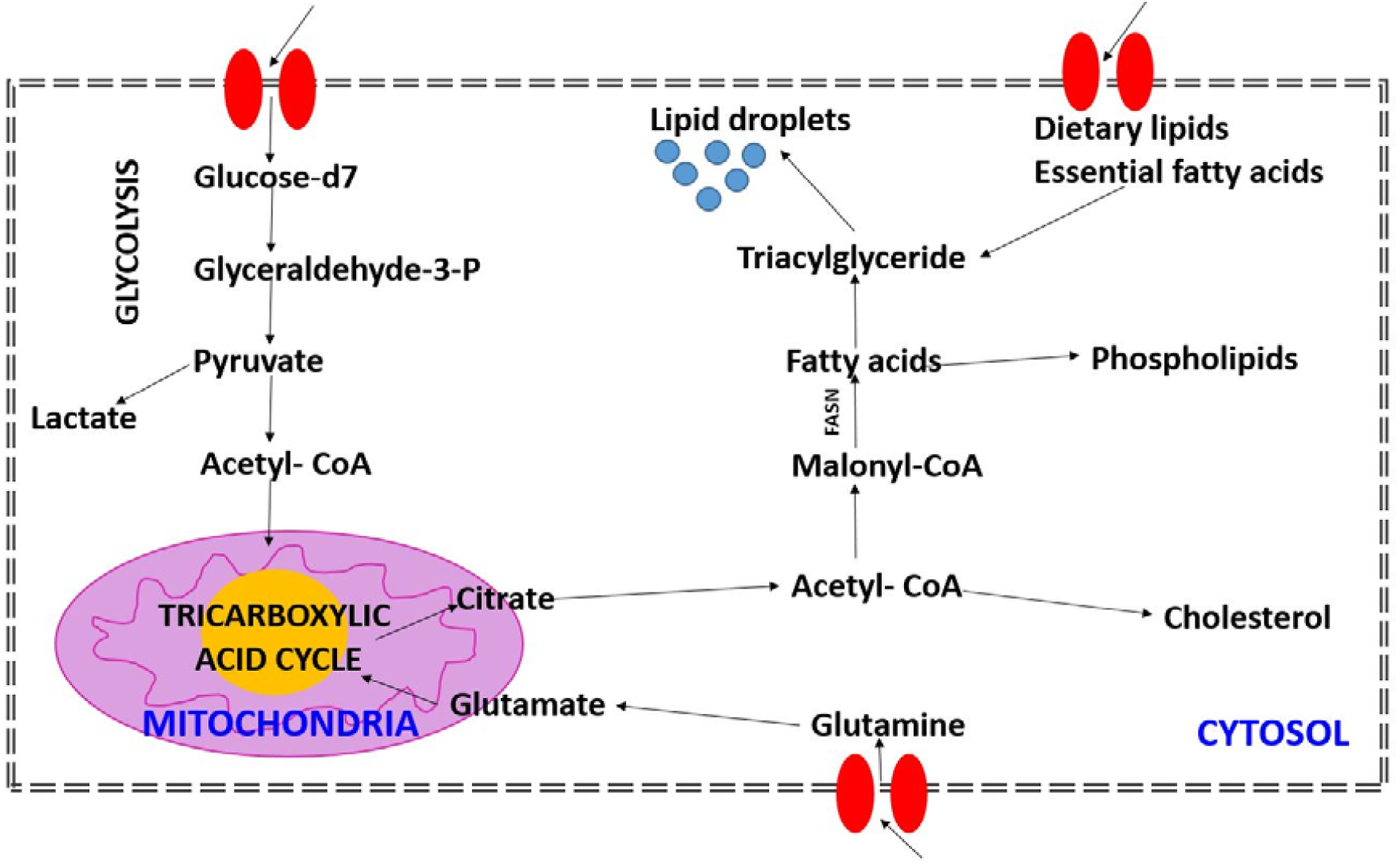
Glucose –derived de-novo lipid synthesis in mammalian cells

Briefly, in normal cells glucose is processed by glycolysis to generate ATP and pyruvate. Then the ribose 5-phosphate and NADPH were produced through the pentose phosphate pathway (PPP), or enter into the tricarboxylic acid (TCA) cycle in mitochondrion. Glucose-derived citrate is converted to acetyl-CoA, oxaloacetate (OAA), or a-ketoglutarate (a-KG). Glutamine is deaminated to form glutamate, which is processed to produce a-KG for use in the TCA cycle.

In contrast, the main pathway of glucose metabolism in cancer cells is aerobic glycolysis, termed Warburg effect. In cancer cells, glucose uptake and the production of lactate was reported to dramatically increase even in the presence of oxygen and fully functioning mitochondria. This classic type of metabolic change provides fast pathway to produce substrates required for cancer cell proliferation and division, which is involved in tumor growth, metastatic progression and long-term survival. It must be emphasized that both glycolysis and mitochondrial metabolism are crucial to cancer cells in the Warburg Effect. [42]

Our results provide a direct evidence that high level of glucose decreases the metabolism via oxidative phosporylation in mitochondria in cancer cells and shifts the metabolism to glycolysis via Wartburg effect. It is worth to emphasize that the Raman intensity of the C-H bands in the 2800-3000 cm^-1^ spectral range for the glucose free cells and cells supplemented with high hyperglycemia level (100 mM) are practically identical, which means that the mitochondrial metabolism is significantly reduced and the Raman signals in this region come mainly from the lipids uptake provided by FBS.

Now, we will show that the comparison between the results for normal and deuterated glucose is an excellent method to monitor *de novo* lipid synthesis and to separate *de novo* synthesis pathways from exogenous uptake of lipids provided with the diet.

Isotope substitution by deuter was used to shed further light on this issue. The stretching vibrations of C−H bonds in the region of 2800-3000 cm^-1^ on the aliphatic backbone of lipids and proteins are found to be effectively decoupled from other molecular vibrations. Each D substitution in a CH_2_ or CH_3_ group of the backbone produces the removal of C-H bonds (and decrease of the Raman signal in the region of 2800-3000 cm^-1^) and an appearance of the band in the C−D stretching region (2100-2300 cm^-1^).

The results presented in Fig. 5 demonstrate that the most significant changes in the spectra regions of 2100-2300 cm^-1^ and 2800-3000 cm^-1^ upon D substitution are observed only in lipid droplets (Fig.5E). It indicates that glucose is largely utilized for *de novo* lipid synthesis lipid synthesis.

In order to better visualize the changes caused in glucose metabolism we concentrated on lipid droplets. We compared the average spectra characteristic for these lipid structures. The average Raman spectra are based on 16 spectra obtained from Raman imaging of a small regions corresponding to lipid droplets recorded for much longer integration times of 20 s. For each type of supplementation we measured 4 cells. The comparison of average normalized Raman spectra of lipid droplets incubated with glucose and deuterated glucose of 5 mM and 100 mM concentration in medium is presented on Fig 6.

**Fig. 6.**
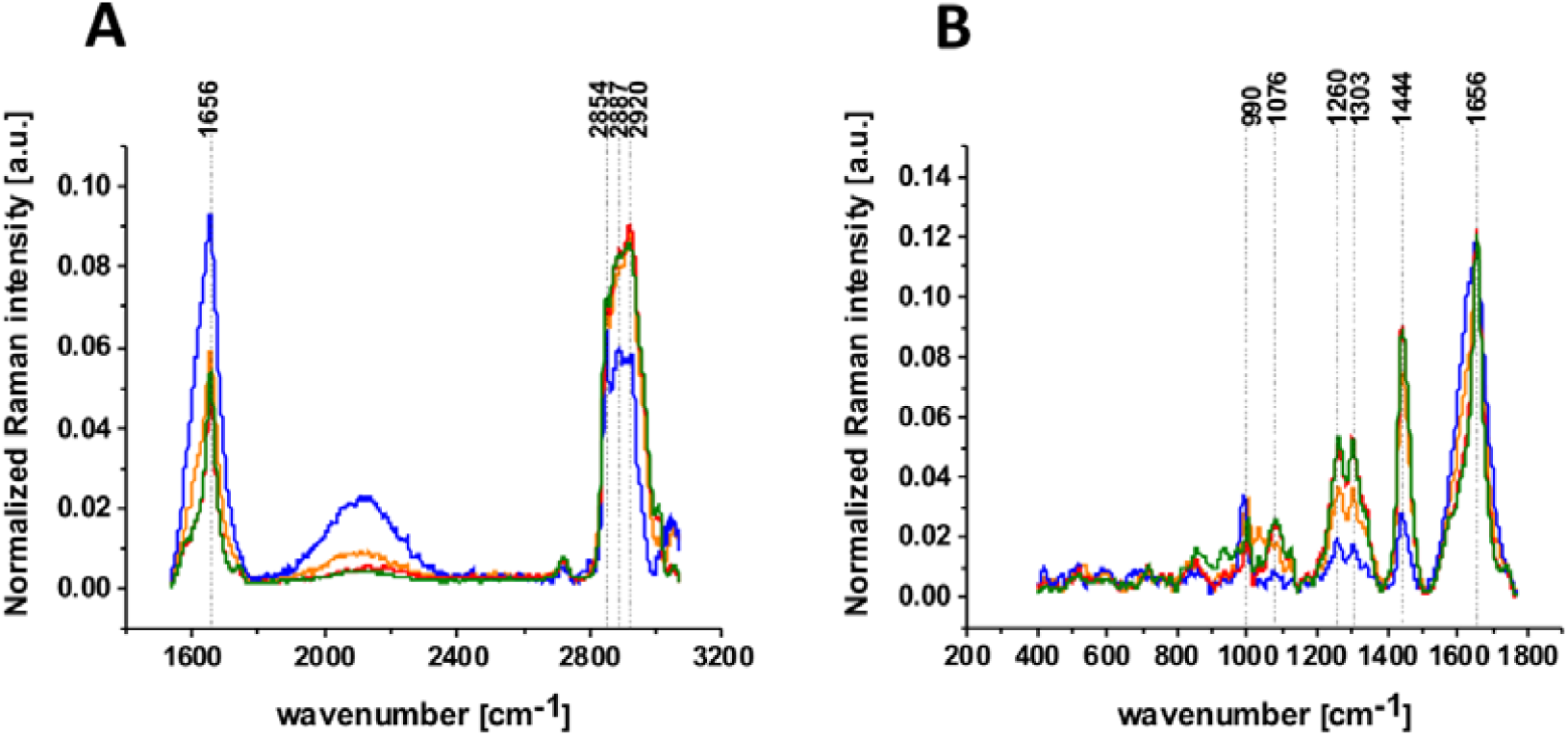
The average Raman spectra for human lung cells for lipid rich area supplemented with glucose 100 mM (orange line), deuterated glucose 100 mM (blue line), glucose 5 mM (green line) and deuterated glucose 5 mM (red line) in high frequency region (A) and in low frequency region (B); integration time 20 s, laser power 10 mW.

One can see from Fig. 6 that the prominent changes are visible in region of amide I and C=C lipid vibrations at 1656 cm^-1^, C-D stretching vibrations of lipids at 2000-2400 cm^-1^ and C-H stretching vibrations of lipids. One can see that the C-D band of lipid droplets gives the strongest Raman signal in cells incubated with 100 mM deuterated glucose while the intensities of bands corresponding to C-H at 2800-3000 cm^-1^ and C=C at 1656 cm^-1^ stretching vibrations decrease due to isotope substitution of C-H bonds by C-D bonds. The results provide direct evidence that glucose is largely utilized for *de novo* lipid synthesis.

The isotope substitution provides information not only on lipid metabolism, but also on another metabolic processes. We could detect other metabolites of glucose by using glucose as precursor for other macromolecular synthesis, such as nucleotides and proteins. Hyperspectral imaging in the fingerprint region would allow us to separate different metabolic species based on the spectra differences.

The results from the fingerprint region presented in Fig. 7 reflect redox mitochondrial metabolic processes.

**Fig. 7.**
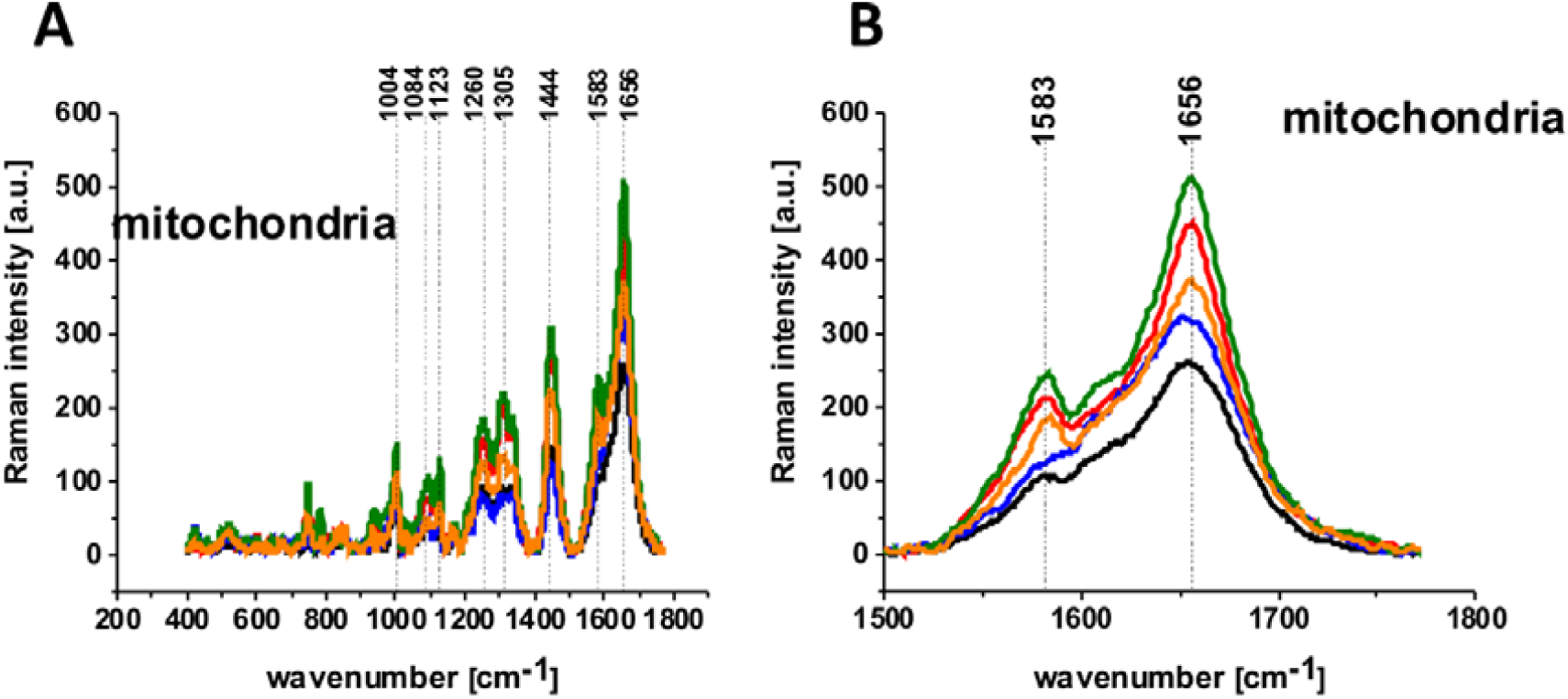
The average Raman spectra of human lung single cell CCL-185 without glucose supplementation (black line), supplemented with normal glucose (green line) and deuterated glucose (red line) at normal conditions (5mM) and supplemented with normal glucose (orange line) and deuterated glucose (blue line) at hyperglycemia conditions (100 mM) for 200-1900 cm^-1^ spectral region (A) and for 1500-1800 cm^-1^ spectral region (B) obtained from Raman images of separate clusters identified by Cluster Analysis method assigned to: mitochondria.

One can see that for normal physiological conditions (5mM) both for normal and deuterated glucose is much the intensity of 1583 cm^-1^ Raman vibration is higher than for hyperglycemia (100 mM).

The band at 1583 cm^-1^ in Fig. 7 represents the “redox state Raman marker” of cytochrome c. Recently we demonstrated that this Raman vibration can serve as a sensitive indicator of oxidized and reduced forms of cytochrome c. [26] It indicates that the Raman peak at 1583 cm^-1^ can be used as a marker to explore apoptosis and oxidative phosphorylation in mitochondria. This band reflects the dual face of cytochrome c in life and death decisions: apoptosis and oxidative phosphorylation. The balance between cancer cells proliferation (oxidative phosphorylation) and death (apoptosis) decide about level of cancer development. [26] [43] The Raman signal of a single cell at 1583 cm^-1^ depends on cytochrome c concentration (which also depends on number of mitochondria in a cell), and the redox state (oxidized or reduced forms). The Raman intensity of the oxidized form is much smaller than that of the reduced form. [34] [26] [44] [25] Inside a normal mitochondrium cytochrome c exists in the oxidized form. Dysfunction of mitochondrium associated with several malignancies, including cancer or hyperglycemia blocks the transfer of electrons between complexes III and IV of the respiratory chain, resulting in decreasing the efficiency of the oxidative phosphorylation (respiration) process and lower ATP synthesis.

It has been suggested that under a hyperglycemic condition, more than usual glucose enters the metabolic cycle. As a result of more intense changes in the tricarboxylic acid cycle, too much NADH and FADH_2_ is supplied to the mitochondrial chain. As a consequence, the voltage gradient across the mitochondrial membrane increases to the critical point where electron transport in complex III is blocked, causing the electrons to be withdrawn back into coenzyme Q where they are attached to oxygen molecules. In this way, superoxide anion radicals are generated. [45]

The intensity of Raman bands at 1444 cm^-1^ corresponding to C-H bending vibrations of lipids in Fig. 7 shows that the signals are lower at hyperglycemia (100 mM of glucose) than for normal level of glucose (5mM).

The results from Fig. 7 support our results obtained from the high frequency region in Fig. 5. It indicates that *de novo* lipid synthesis decreases at hyperglycemia conditions for cancer lung cells.

## Conclusions

This paper presents a truly unique landscape of cancer cells biochemistry by non-invasive Raman method to study glucose metabolism in lung cancer in normal and hyperglycemia conditions. We found that isotope substitution of glucose by deuterated glucose allows to separate *de novo* lipid synthesis from exogenous uptake of lipids obtained from the diet. We demonstrated that glucose is largely utilized for de novo lipid synthesis lipid synthesis. Our results provide a direct evidence that high level of glucose decreases the metabolism via oxidative phosporylation in mitochondria in cancer cells and shifts the metabolism to glycolysis via Wartburg effect.

## Acknowledgement

This work was supported by the National Science Centre of Poland (Narodowe Centrum Nauki, UMO-2019/33/B/ST4/01961) and “FU^2^N-Fund for the Improvement of the Skills of Young Scientists” supporting scientific excellence of the Lodz University of Technology-grant no. W3/2P/2022.

