## Supplemental Data 1 for "Hyperglycemia and Cancer. Human lung carcinoma by means of Raman spectroscopy and imaging": supplementary materials.docx

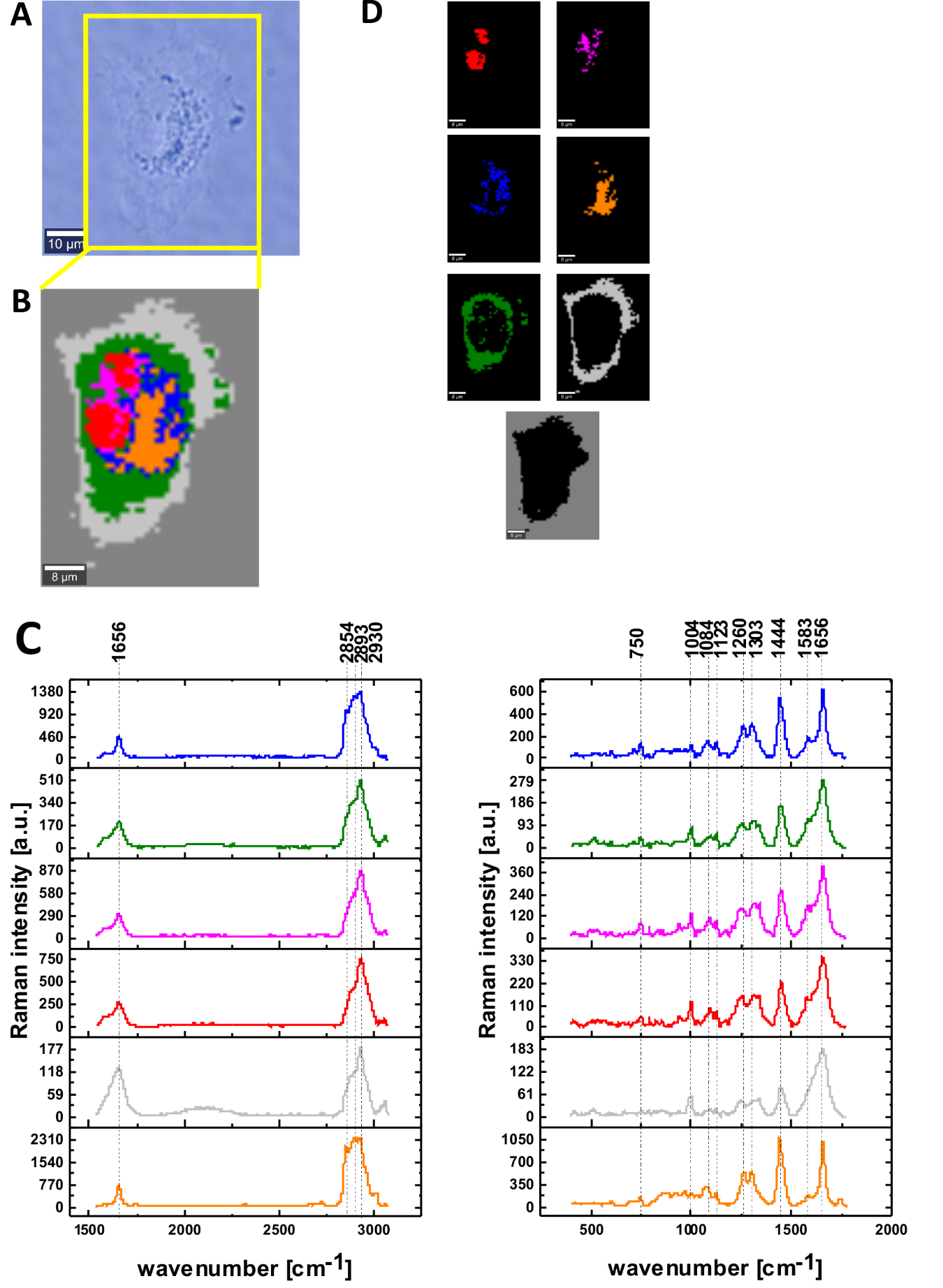

*Fig SM1 The microscopy image (A), Raman image for area marked by yellow frame in the panel A, the size of Raman image (44 μm × 60 μm), resolution 1 μm of human lung single cell CCL-185 supplemented with glucose 5 mM constructed based on Cluster Analysis method (B), the average Raman spectra for all clusters for high and for low frequency region (C), Raman images of separate clusters identified by Cluster Analysis method assigned to: nucleus (red), mitochondria (magenta), lipid regions (blue and orange), cytoplasm (green), cell membrane (light grey) and cell environment (dark grey) (D) colors of the spectra correspond to the colors of clusters; integration time 0.3 sec in the high frequency region and 0.5 sec in the fingerprint region, laser power 10mW.*

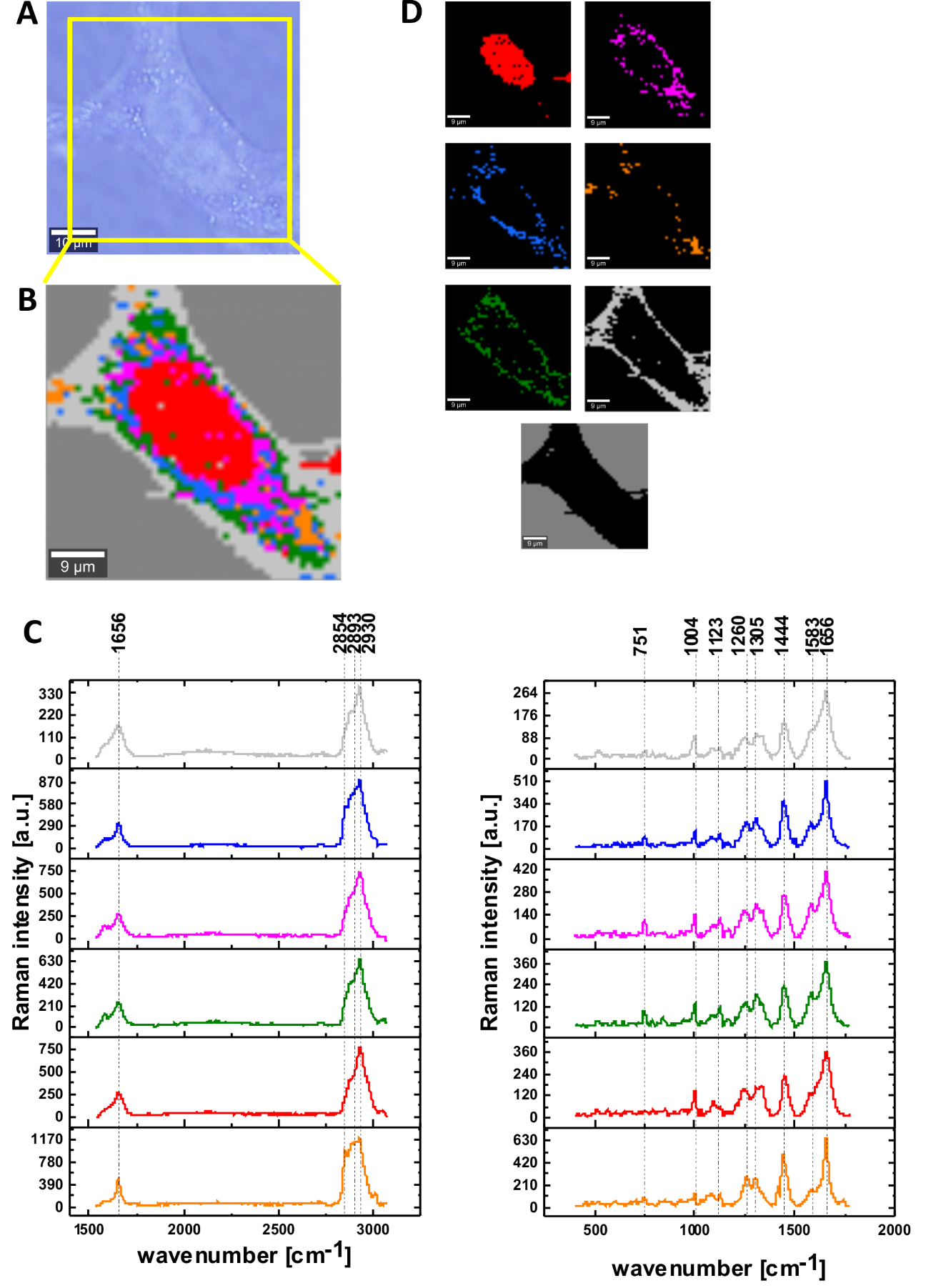

*Fig SM2 The microscopy image (A), Raman image for area marked by yellow frame in the panel A, the size of Raman image (50 μm × 50 μm), resolution 1 μm of human lung single cell CCL-185 supplemented with deuterated glucose 5 mM constructed based on Cluster Analysis method (B), the average Raman spectra for all clusters for high and for low frequency region (C), Raman images of separate clusters identified by Cluster Analysis method assigned to: nucleus (red), mitochondria (magenta), lipid regions (blue and orange), cytoplasm (green), cell membrane (light grey) and cell environment (dark grey) (D) colors of the spectra correspond to the colors of clusters; integration time 0.3 sec in the high frequency region and 0.5 sec in the fingerprint region, laser power 10mW.*
